# Reconstructing the dynamics of past coral endosymbiotic algae communities using coral ancient DNA (*cora*DNA)

**DOI:** 10.1101/2024.03.21.584420

**Authors:** Olivier Rey, Delphine Dissard, Eve Toulza, Thomas Guinebert, Mathilde Saccas, Jean-François Allienne, John Butsher, Mourad BenSalah Zoubir, G. Iwankow, Christelle Tougard, Jérémie Vidal-Dupiol

## Abstract

Most scleractinian corals are drastically threatened due to global changes but some colonies are intriguingly resistant to heat stress. Coral thermal tolerance partly relies on genomic determinism among the cnidarian compartment but also on the physiology of their associated symbiotic algae (Symbiodiniaceae). In fact, some corals can shift and/or shuffle their associated Symbiodiniaceae communities to temporally cope with heat stress. So far coral adjustments of their endosymbiotic algae were mainly observed at short-term evolutionary time scales and we lack a general vision of coral holobiont evolution at broader timescales. We here combined the use of ancient DNA from a coral core and a metabarcoding approach, to retrace past Symbiodiniaceae communities associated with a living colony of *Porites lobata* from New Caledonia over the last century. We were able to extract ancient DNA along the coral core at 19 time points dating back to the 1870’s. Overall, we detected 13 OTUs, nine of which were affiliated to the Symbiodiniaceae *Cladocopium* clade, one to *Azadinium spinosum* (Dinophycae); one to the host *P. lobata*, the two other OTUs remained unidentified. One OTU was largely predominant and was ubiquitous over all samples. The number of OTUs was marginally correlated to the total number of sequences per sample but not to the age of the *cora*DNA sample. We found a generally stable core microbiota associated with *P. lobata*, although drastic change in community composition was observed in coraDNA samples corresponding to an extreme hot winter temperature event. More generally, this study paves the way for further investigations on the evolutionary dynamics of coral holobionts at the colony level over large temporal scales.

## Introduction

Scleractinian corals and the associated reef ecosystems are among the most threatened systems worldwide due to global changes. Almost half of all living corals have already been destroyed in the last 150 years ^1^. Such massive loss has accelerated over the last three decades under the influence of extreme climatic events, in particular heat waves, the frequency of which is steadily increasing ^1^. Despite this alarming situation, coral colonies particularly resistant to heat stress have been identified ^2–4^. These observations provide some hope for the maintenance and/ or the restoration of corals and ecosystems they support. They also call for an urging need to unravel the molecular mechanisms by which some coral colonies survive through time despite recurrent events of environmental stresses and help adjust conservation plans. This task is all but trivial in particular because corals are complex holobionts composed of cnidarians associated with microbial communities including – among others – endosymbiotic algae (Symbiodiniaceae), protista and bacteria, each of these partners potentially influencing the thermotolerance of the colonies ^5,6^.

Variation in coral thermal tolerance across latitude, at least partly relies on genomic variation among the cnidarian compartment of coral colonies ^7^. In this respect, it has been suggested that resistant genotypes could emerge through intrinsic rapid genomic changes such as somatic mutations ^8^, the activation of transposable elements ^9^ and/ or some modifications in the proportions of genotypes coexisting within the same colony ^10,11^. Additionally to genomic modifications, changes in DNA methylation patterns in cnidarians may also induce – at least temporarily – adaptive phenotypic adjustments in coral colonies exposed to recurrent heat stress ^12,13^. Besides changes within the cnidarian compartments, coral thermal tolerance also depends on the physiology of their associated symbiotic algae (Symbiodiniaceae), with which they form a phototrophic mutualistic symbiosis ^14,15^. Symbiodiniaceae constitute a highly diversified taxonomic group among which the identified species and even strains among species display huge variation in thermal tolerance ^16,17^. Recently, a laboratory experiment has shown that experimentally adapted strains of the Symbiodiniaceae *Cladocopium goreaui* to high temperature, provide a better protection to heat stress in *Acropora tenuis* colonies after reimplantation ^18^. This suggests that thermal tolerance may be acquired rapidly by natural coral colonies *via* the acquisition of acclimatized or adapted Symbiodiniaceae from the surrounding environment in response to heat stress. In fact, symbiont switching – the acquisition of new (thermally resistant) species/strain from the environment – and symbiont shuffling – the modification of the relative abundance of the inner Symbiodiniaceae strains within host – constitute two alternative rapid responses for corals to temporarily cope with heat waves ^19,20^. Finally, while other partners involved within the coral holobiont such as bacteria, fungi, viruses and protists, certainly display higher adaptive capacity than corals in particular due to their short generation time, their role in fostering adaptive response to heat stress at the holobiont scale still remain to be demonstrated ^5,13,21^.

Most of our knowledge on the mechanisms underlying the adaptive response of coral colonies to global changes relies on empirical studies based on controlled experiments conducted either in laboratory or in the field, over extremely short-term evolutionary time scales. These approaches are crucial to dissect the relative importance of specific molecular mechanisms in action to foster coral responses. They have dramatically changed our vision on the short-term adaptive capacity of corals ^12,13^. However, and complementary to these approaches, we need a more general vision of coral holobiont evolution at broader timescales to assess their evolutionary dynamics in response to the recent fast evolving environmental changes. In this respect, ancient DNA (aDNA) based approaches are promising ^22^. These approaches generally rely on DNA extracted from archaeological or paleontological remains or from museum samples. For instance, museum samples of 8 octocoral species originally collected from successive epochs revealed that mutualism between several coral species and their associated Symbiodiniaceae remained stable overtime with no major changes in the last two centuries despite major anthropogenic global change ^23^. In a more general context of biodiversity, aDNA was also applied to reconstruct the community assemblages of reef ecosystems from sediment cores ^24,25^. Surprisingly however, no attempts were made to extract aDNA directly from cores excavated from a living massive coral colony that chronicles decades and up to centuries, of its lifetime. Such an approach is particularly promising because it could allow accessing the DNA material from individual holobiontic colonies over time and thus reconstructing their intrinsic eco-evolutionary history.

We here provide the first proof of concept of the use of aDNA from a coral core, hereafter called *cora*DNA (referring to the recent *seda*DNA approach developed to study DNA from sedimentary cores), to reconstruct past Symbiodiniaceae communities associated with a living colony of *Porites lobata* over the last century. We discuss the benefit of such approach to unravel the eco-evolutionary dynamics of coral holobiont and the current technical limitations that will need to be bypassed to have access to DNA from the cnidarian compartment and hence reconstruct the full eco-evolutionary trajectories of coral holobionts over time.

## Material and methods

### Core sampling

*Porites lobata* is a massive reef-building scleractinian coral ubiquitous over the Tropical Pacific Ocean. Colonies can be extremely long-lived (up to several centuries) and their growth strongly depend on the environmental conditions in which they grow, especially seawater temperature ^26^. These characteristics make this species particularly suitable in the context of this study since it makes it possible to obtain long cores and the marked seasonal successions of growth allow the collection of core matrix at the year scale from which DNA can be extracted. A coral-matrix core was collected from a 2.5-m high colony of *Porites lobata* at 10-m depth within the Southern Lagoon of New Caledonia (22°17,146 et 166°11,004) the 17^th^ of December 2018 (Figure 1A..). The excavated 8-cm diameter and ∼ 80-cm long core was immediately stored in dry ice and transferred under freezing conditions to the Institute de Recherche pour le Developpement (IRD) in Nouméa, prior being sent in dry ice to the IHPE laboratory in Perpignan where it was stored at -80°C until subsequent analyses. Thus, the core has been kept at -80°C from its initial excavation until the sampling of *cora*DNA.

**Figure 1:**
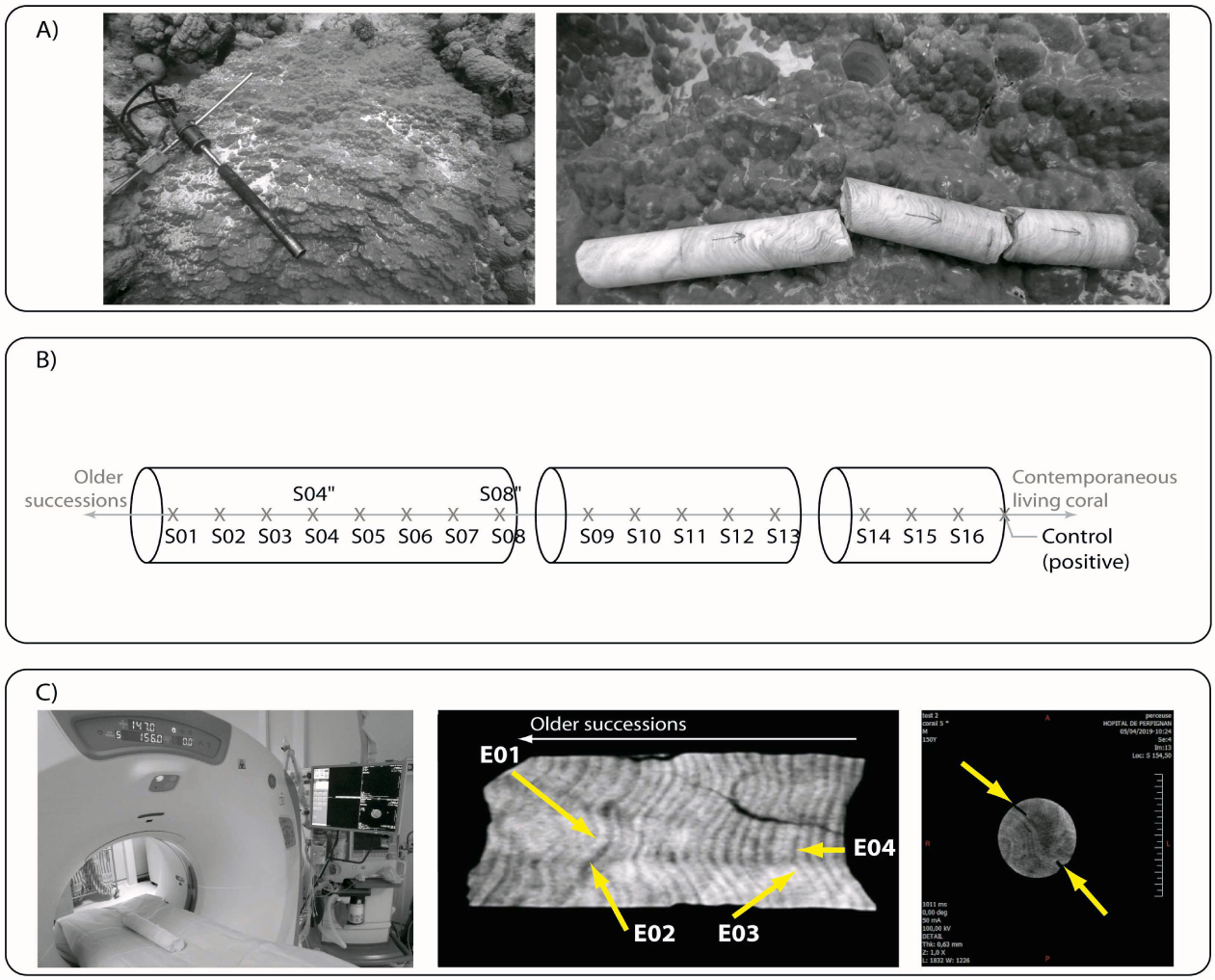
Details on the coral core of the studied Porites lobata colony. Pictures of its excavation in December 2018 from the living colony in New Caledonia are provided in (A). In (B), the coraDNA samples that were extracted along the core are represented from the older sample (S01) to the contemporaneous living coral (control). In (C) Pictures of the interventional sampling using a CT scan of the four coraDNA samples corresponding to successions before (E01) and after (E02) the 1997-1998 ENSO event; and before (E03) and after (E04) the anormal hot winter in 2010. Yellow arrows in the last panel in (C) point toward the hole left by the drilling bit after the sampling of the E04 coraDNA sample.

### Dating successions and sampling ancient DNA from the coral core

The aging of the coral core was determined based on the counting of density bands as described in Wu *et al.* ^27^. Annual skeletal density bands were observed using a CT scan at the imagery service of the public Hospital of Perpignan (France). The observed density bands result from variation in colony growth rate during the winter (slow) and during the summer (rapid) that lead to a more or less dense aragonite skeleton along the year ^26^. According to this pattern, one year was reconstructed by summing a clear (summer) and dark (winter) density bands ^27^. Based on the reconstructed chronology of the sampled coral colony, a total of 23 samples were collected along the coral core (Figure 1.B and C). Briefly, each sampling location along the core was first rapidly washed using a 10% NaClO solution and ∼ 3 mm of the external layer was removed from the core surface using a sterile disposable scalpel to avoid possible DNA contamination with contemporaneous genomic material. Once the surface of the core was decontaminated, an electric driller with an individualized sterile 2-mm diameter drill bit was used to extract core powder along a ∼ 1.5 – 2 cm depth drilling hole perpendicular to the surface of the core. The bits containing the powder was immediately transferred and rinsed in a 2 ml Eppendorf (DNA/RNA and DNase/RNase free tube) containing 1 ml of Tissue Lysis buffer ATL from the QIAamp DNA Micro Kit (Qiagen) and stored at 4°C until the DNA extraction process (i.e. at most 24 h later).

Sampling along the core was achieved following two strategies. The first strategy consisted in systematically sampling every 5 cm along the core without accurate dating. According to the retraced chronology and to the thickness of the observed density bands, such strategy roughly corresponded to a sample collected every 8 to 10 years back in time over the colony history from December 2018 (collection date) until the 1870’s. A total of 17 *cora*DNA samples were extracted following this design including one sample from the top of the coral core (as positive control) which corresponds to the living colony at the time of the core sampling (Sample S01 to S16 and control in Figure 1.B.). This strategy was used to specifically test our ability to detect and obtain processable *cora*DNA samples over large timescales. Moreover, two duplicates were sampled for two random samples (S04 and S08; Figure 1B.). These duplicates consisted in independent *cora*DNA samples obtained from the same section along the core and using two independent sterile drill bits.

The second strategy aimed at studying the dynamics of the algae endosymbiotic communities associated with coral colonies during well documented past extreme climatic events. We targeted two well-known climatic anomalies that were easily observable along the core in 2010 (abnormally hot winter) and in 1997-1998 which corresponded to a severe ENSO event ^27^. Two samples were collected, one prior and the other after each of these two climatic events (N = 4; Figure 1C.). For this purpose, and to accurately sample the targeted coral successions, the drilling procedure was achieved under interventional CT 3D scan at the imagery services from the public Hospital of Perpignan. The coordinates of the targeted successions were first retrieved, and the drilling sampling was achieved under interventional scan to ensure that the same succession was sampled within the core during the sampling process (Figure 1C.). The precautions previously described were taken to avoid contamination (treatment with bleached plus abrasion of the ∼3 mm of the external layer of the core and a unique sterile drill bit per sample).

Additionally to the overall 23 samples collected along the core, a negative control was prepared at the time of the sampling collection to check for possible contaminations over the whole process from DNA sampling to library preparation. This negative control was obtained following the same protocol as described above except that the drill bit did not touch the core prior to the rinsing step into the Tissue Lysis buffer ATL.

### DNA extraction

The overall DNA extraction process and DNA amplification steps were performed at the degraded DNA platform (Institut des Sciences de l’Evolution de Montpellier, France; http://club.quomodo.com/plateforme-adn-degrade) offering facilities dedicated to the study of ancient DNA. DNA extractions were performed using the QIAamp DNA Micro Kit (Qiagen) following the “*Purification of genomic DNA from bones*” protocol. Briefly, 25 µl of Qiagen Proteinase K were added to each sample. Samples were then incubated at 56°C with shaking at 1200 rpm overnight. The day after, 1 ml of AL buffer was added and samples were incubated at 70°C during 10 minutes to inactivate enzymes. After a centrifugation step at full speed during one minute, the supernatant was transferred to a QIAamp MinElute column. Once the DNA attached on the column membrane (centrifugation step at 8000 rpm for 1 minute), the membrane was successively washed using 600 µl of the AW1 and the AW2 buffer. After these washing steps, the membrane was dried by centrifugation during 3 minutes at full speed. The DNA was eluted using 30 µL of sterile water. The eluted DNA samples were then stored at -20°C until subsequent molecular processes. The quality of DNA extracts wase assessed on a Bioanalyzer High Sensitivity DNA kit (Agilent, USA) and DNA was quantified using a Qubit fluorometric quantification with the ds DNA High Sensitivity assay kit (Thermo Fisher Scientific, USA).

### Library preparation and sequencing

Each DNA sample was used as initial template for amplifying a portion of the *ITS2* gene using a set of 4 primers derived from the literature ^28,29^ with Illumina adapters and spacers (Table 1).

**Table 1:**
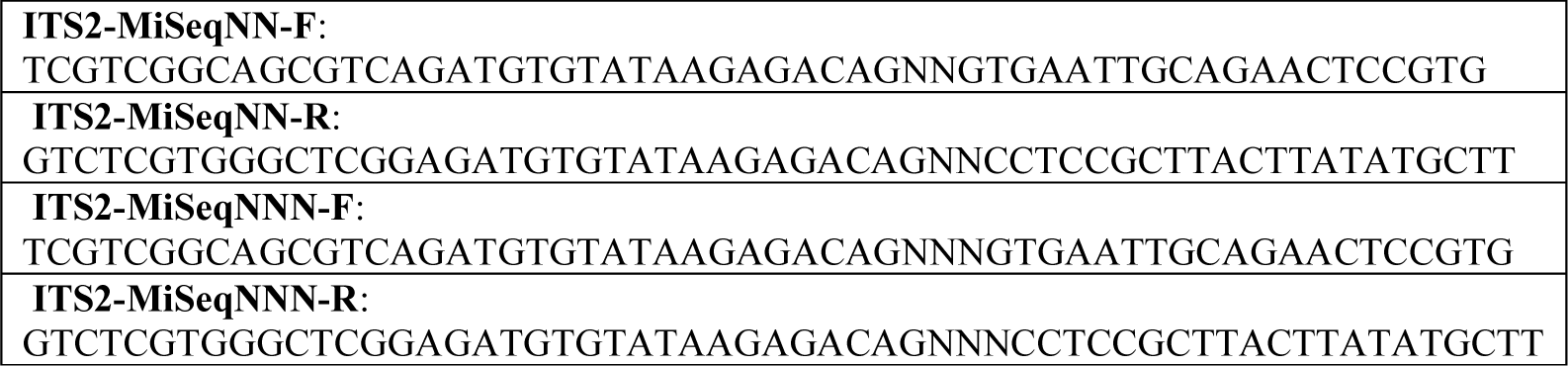
Sequence of the 4 primers (including adapters and spacers) used to amplify a ∼ 350 bp fragment of the ITS2 gene.

Concomitantly, the extraction negative control and a PCR negative control (consisting in distilled water) were used in the same PCR reaction to control for potential contamination during the overall process. PCRs were performed in 50 µL containing 5µL of Buffer 10X PCR Gold (1X), 5 µL of MgCL2 (25 mM), 1 µL of each dNTPs (200 µM), 1.35 µL of primer mix (0.25 µM), 0.25 µL of Taq Polymerase (1.25 units), 4 µL of DNA template and 35.4 µL of distilled sterile water. Amplification was performed using the following PCR conditions: an initial denaturation step at 95°C during 7 minutes followed by 35 cycles each of which constituted of a denaturation step at 95°C during 15 sec, an annealing step at 57°C during 30 sec and an elongation step at 72°C during 30 sec. After these 35 cycles, a final elongation step at 72°C during 7 minutes was applied. The amplification was checked by migrating 5 µL of each PCR reaction on a 2% agarose gel stained with ethidium bromide and visualized under UVs. Expected size of the amplicon is around 350 bp.

Indexed libraries were generated using the standard Illumina two-step PCR protocol using Q5 high fidelity DNA polymerase (New England Biolabs). Paired-end sequencing with a 2×250 bp read length was performed at the Bio-Environment platform (University of Perpignan Via Domitia Perpignan, France) on a MiSeq system (Illumina) using v2 chemistry according to the manufacturer’s protocol. Sequencing data are available for download on SRA under the bioproject number ###

### Data processing

The sequence datasets were uploaded to the Galaxy web platform ^30^ and processed using the Finding Rapidly OTUs with Galaxy Solution (FROGS) pipeline at the GenoToul platform (Toulouse, France) ^31^. The first pre-processing step of this pipeline consisted in demultiplexing, dereplicating and cleaning all reads. Given the theoretical expected amplicon size (i.e., ∼ 350 bp) and after a quick overview of the overall amplicon lengths, we kept all sequences which sizes ranged from 150 to 490 pb. These filtered sequences were then clustered using the SWARM algorithm using an aggregation distance set at 1. This iterative clustering approach uses amplicons’ homology, structure and abundance hence limiting potential biases generated by other approaches such as *de novo* clustering methods (i.e., input order dependencies and arbitrary clustering) (Mahe et al., 2014). Clusters were then cleaned to remove potential chimeras, singletons and under-represented clusters (i.e. <50 sequences) using VSEARCH (Rognes et al., 2016).

Each OTU was identified based on a nucleotide megablast using the online standard database (nt/nr) available from NCBI. To more precisely identify OTUs that were affiliated to the Symbiodiniaceae genus *Cladocopium* (clade C) based on the results from Blast, we next computed pairwise genetic distance between each OTU seed sequence and a subset of sequences from different *Cladocopium* strains available from LaJeunesse *et al.* ^32^. Pairwise genetic distances were computed using the ‘K80’ evolutionary model as implemented in the ape package V5.0 in R ^33^.

## Results

### Dating back coral successions along the core

The obtained coral-skeleton core followed the colony’s growth axis on more than 2/3 of its length (see Figure 1C.; middle panel for illustration). We were hence capable to accurately date back the growth successions over the last seven decades, i.e., back to 1950. Density bands along the oldest part of the coral core partly deviated from the central vertical axis preventing our capability to date back accurately the oldest successions. Annual growth bands were approximately 5-6 mm thick and generally homogenous along the core. According to this measure, we estimated that the oldest *cora*DNA sample collected from the core (S01; Figure 1B.) roughly correspond to coral holobiont that lived in the 1870’s.

### Amplicon sequencing from coraDNA samples, sequence affiliation and identification of endosymbiotic algae communities

Because of low DNA concentrations, DNA quality and quantity could not be assessed from all samples except the positive control. PCR amplicons of expected size for ITS2 Symbiodiniaceae marker were however obtained from all samples. After sequencing and filtration steps, sequences were validated from 19 out of the 23 initial *cora*DNA samples including the positive control (Table 2). The final number of filtered sequences obtained from these 19 *cora*DNA samples ranged from 1476 (sample S12) to 76403 (sample S13) (Table 2). Four *cora*DNA samples displayed negligible numbers of sequences (from 9 to 33) and were then excluded for further analyses. Similarly, extraction and PCR negative controls exhibited negligible numbers of sequences (8 and 17, respectively) with an expected size of 300 bp (190 – 450 bp; Table 1), which indicates negligible cross-contaminations during all the process from sampling, extraction to sequencing.

**Table 2:** Details on the obtained and conserved sequences from each coraDNA sample.

| Sample | % kept | Paired-end assembled (%) | With 5' primer (%) | With 3' primer (%) | With expected length (%) | Without Ns (%) | Nseq OTU > 50 seq |
| --- | --- | --- | --- | --- | --- | --- | --- |
| **<u>Samples kept in the analyses</u>** |  |  |  |  |  |  |  |
| E01 | 97.29 | 77.44 | 77.38 | 76.15 | 75.34 | 75.34 | 73241 |
| E02 | 97.12 | 57.91 | 57.87 | 56.68 | 56.24 | 56.24 | 54571 |
| E03 | 96.37 | 73.18 | 73.13 | 71.18 | 70.52 | 70.52 | 68484 |
| E04 | 91.60 | 48.53 | 48.52 | 47.35 | 44.46 | 44.46 | 43454 |
| S01 | 87.58 | 75.61 | 75.58 | 74.20 | 66.21 | 66.21 | 64448 |
| S04 | 75.10 | 59.92 | 59.90 | 59.09 | 45.00 | 45.00 | 44203 |
| S04" | 85.18 | 15.65 | 15.62 | 15.30 | 13.33 | 13.33 | 12993 |
| S05 | 4.44 | 89.98 | 89.96 | 89.83 | 3.99 | 3.99 | 3928 |
| S06 | 67.49 | 65.52 | 65.50 | 63.76 | 44.22 | 44.22 | 42583 |
| S08 | 96.01 | 77.93 | 77.90 | 75.11 | 74.82 | 74.82 | 71366 |
| S08" | 97.56 | 46.68 | 46.63 | 45.66 | 45.54 | 45.54 | 44501 |
| S09 | 63.04 | 70.01 | 69.98 | 68.80 | 44.13 | 44.13 | 43147 |
| S10 | 8.32 | 48.20 | 48.19 | 48.09 | 4.012 | 4.012 | 3884 |
| S11 | 85.80 | 54.13 | 54.11 | 53.04 | 46.44 | 46.44 | 45236 |
| S12 | 6.98 | 21.95 | 21.94 | 21.90 | 1.53 | 1.53 | 1476 |
| S13 | 96.83 | 80.93 | 80.90 | 78.79 | 78.36 | 78.36 | 76403 |
| S14 | 93.24 | 74.74 | 74.69 | 73.05 | 69.69 | 69.69 | 68031 |
| S15 | 56.67 | 88.20 | 88.18 | 86.22 | 49.98 | 49.98 | 48388 |
| Control | 28.60 | 8.73 | 8.73 | 8.66 | 2.50 | 2.50 | 2454 |
| **<u>Samples removed from the analyses</u>** |  |  |  |  |  |  |  |
| S02 | 0.02 | 152.242 | 152.221 | 152.075 | 33 | 33 | 0 |
| S03 | 0.02 | 58.41 | 58.406 | 58.344 | 9 | 9 | 0 |
| S07 | 0.02 | 76.544 | 76.531 | 76.462 | 15 | 15 | 0 |
| S16 | 0.02 | 52.119 | 52.104 | 52.046 | 11 | 11 | 0 |
| **<u>Negative controls</u>** |  |  |  |  |  |  |  |
| Extraction | 0.02 | 45.646 | 45.644 | 45.597 | 8 | 8 | 0 |
| PCR | 0.15 | 11.01 | 11.007 | 10.998 | 17 | 17 | 0 |

The sequences clustered into 13 OTUs. Nine of these OTUS were affiliated to Symbiodiniaceae and all of which were assigned to the *Cladocopium* clade. Based on the genetic distances computed between each of these 9 clusters and *ITS2* sequences from LaJeunesse *et al*. ^32^, all OTUs were more specifically affiliated to the *ITS2*-C15 type (computed distances ranged from 0 to 0.044; Table S1). Among the 4 other non-specific OTUs, one was affiliated to *Azadinium spinosum* (Dinophycae; OTU_7; Query cover = 100%, E-Value = 6e^-156^; Per. Ident. = 98.1%); one to *P. lobata* (the coral host species; OTU_11; Query cover = 100%, E-Value = 8e^-175^; Per. Ident. = 100%), and the two others were not successfully affiliated with any identified sequence available from the NCBI nr database (OTU_12 and OTU_13). Sequences of *Azadinium spinosum* (OTU_7) were amplified only in the S08 and its duplicate S08” *cora*DNA samples. Amplified sequences of *P. lobata* (OTU_11) were found in the S01, S08, E02 and E03 *cora*DNA samples although at low abundance (Figure 2; Table S2). Finally, OTUs that were not affiliated with any organism documented in the NCBI database (OTU_12 and OTU_13) were present in samples S08 and/or S08” again with low read numbers (Figure 2).

**Figure 2:**
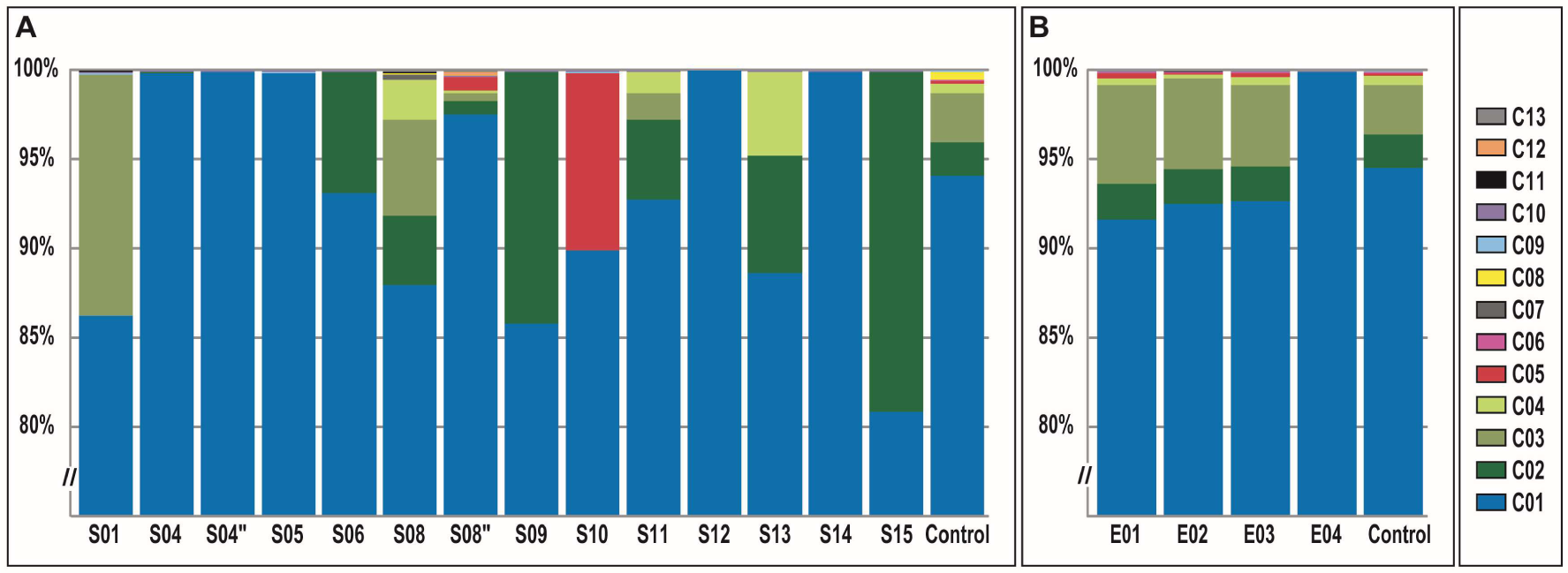
Proportion of sequences attributed to each of the 13 clusters among the 19 positive *cora*DNA samples A. obtained from the systematic sampling (i.e. every 5 cm along the coral core) and B. collected under CT SCAN corresponding to successions before (E01) and after (E02) the 1997-1998 ENSO event; and before (E03) and after (E04) the anormal hot winter in 2010.

### Distribution of Symbiodinium OTUs over the coraDNA samples

The total number of OTUs obtained from each *cora*DNA sample varied from 1 to 10. Among the 9 Symbiodiniaceae OTUs identified, 1 OTU (OTU_1, Figure 2) was clearly predominant representing 92% of the generated sequences overall samples (from 80.88 % in S15 to 100 % in S12). Only OTU_1 is ubiquitous among all *cora*DNA samples. OTU_9 and OTU_10 could also be reasonably considered as ubiquitous since they have been detected in all except the S12 and the positive control *cora*DNA samples. However, these two samples were the one harboring the lowest sequence coverage which may explain this absence (Table 1).

The number of OTUs per sample is marginally correlated to the total number of sequences obtained per sample (Spearman correlation, rho = 0.53; p-value = 0.04) but is not correlated to the age of the *cora*DNA sample (using the position order at which the *cora*DNA was sampled along the core as a proxy; rho = 0.148; p-value = 0.60).

## Discussion

### Unlocking the access to past coral communities

According to our results, ancient DNA from a coral core (*cora*DNA) excavated from living *Porites lobata* colonies can be obtained and used for metabarcoding approaches to reconstruct the Symbiodiniaceae endosymbiotic communities that have been successively associated to the colonies over a century. This paves the way for unprecedented time-series studies to document the eco-evolutionary dynamics ofthe coral/Symbiodiniaceae mutualistic interaction to most massive coral species.

Nine of the 13 OTUs detected in this study belong to the Symbiodiniaceae and all were affiliated to the *ITS2*-C15 type. Since pairwise genetic distances computed between each of these OTUs were relatively low, it is likely that the observed diversity among these strains result from intragenomic variation at the *ITS2* marker as previously highlighted ^34,35^. Importantly however, the possible resulting overestimation of the diversity in the Symbiodiniaceae community documented here cannot be related to possible post-mortem DNA modifications commonly observed in ancient DNA samples ^36^. Indeed, all but one OTU were detected in the living part of the coral core (positive control) which means that the nature and state of the *cora*DNA samples does not seem to artificially inflate diversity of the Symbiodiniaceae community.

Moreover, among the 13 identified OTUs, 1 was attributed to *P. lobata*. This *P. lobata* OTU was detected, although at low abundance, in 4 *cora*DNA samples that do not have obvious commonalities (e.g. spatial location along the core) but a high number of sequences. The detection of the coral host species using the present *ITS2* metabarcoding confirms that host DNA may also be extracted and amplified to some extent. This result makes it possible to consider the exploitation of *cora*DNA samples to trace the eco-evolutionary dynamics of the genome and/or epigenome of the host coral species *P. lobata* at the colony scale.

Among the OTUs not affiliated to Symbiodiniaceae or to the cnidarian, we unexpectedly detected *A. spinosum* among two *cora*DNA duplicates collected from the same location along the core. These duplicates correspond to coral holobionts that lived in the 1940’s – 1950’s. *Azadinium spinosum* is a photosynthetic dinoflagellate that was first described in the late 2000’s and since then, has been reported on the coasts of Northern Europe ^37^, South and Central America ^38,39^ and very recently in Asian pacific ^40^. While we found no direct evidence of its occurrence in New Caledonia or the Southern Pacific exists in the literature, *Azadinium spp.* was recently observed in French Polynesia under a scanning electron microscope (Mirielle Chanin; pers. comm). Importantly *A. spinosum* is one of the primary producer of azaspiracid toxins causing important health issues to several animals, mainly vertebrates including humans, through toxin bioaccumulation ^41^. In this context, azaspiracid-2 (AZA2) toxins were recently detected in New Caledonia based on a SPATT (Solid Phase Adsorption Toxin Tracking) approach, hence indirectly indicating that *Azadinium spp.* can be present locally at least temporary ^42^. So far, no interactions were described between *P. lobata* or any other coral species and *A. spinosum*. We here hypothesize that *A. spinosum* could have been accidentally captured by coral hosts during feeding. Accordingly, the detection of *A. spinosum* at only one time period (although detected in two *cora*DNA duplicate samples from this time period) could coincide with a *A. spinosum* bloom episode as previously described at some localities ^38^. Thus, additionally to DNA from Symbiodiniaceae and their coral hosts, our results suggest that we can also detect the punctual presence back in time of other organisms (most likely present in high abundance) in the environment and that may have been captured by corals. However, (short) specific molecular markers tailored to the targeted organisms would be necessary, as the marker used in this study was not initially designed for such purposes.

From a technical point of view, it is important to note that the number of OTUs detected in the *cora*DNA samples was associated to sequencing output but not to the age of the *cora*DNA samples (estimated from the position of the *cora*DNA samples along the core). This suggests that a low number of sequences but probably not the age of the *cora*DNA could lead to an under represented vision of the studied past Symbiodiniaceae communities (see also rarefaction curves in Supplementary Figure SF1). This is clearly illustrated by the S12 *cora*DNA fairly recent sample, for which we obtained the smallest number of sequences and that harbors only one OTU (OTU_1; Figure 2) which is also predominant in all *cora*DNA samples (representing 80.9 % to 100 % percent of the overall filtered sequences). Conversely, we detected up to 5 different clusters in the oldest *cora*DNA sample (S01) including 4 OTUs affiliated to *Cladocopium* and one affiliated to *P. lobata*. We thus advise to sequence libraries from *cora*DNA samples with a high coverage to avoid possible false negatives or to use a rarefaction approach to characterize the optimal sequencing depth.

### Dynamics of Symbiodiniaceae at large temporal scale

In addition to breaking down a technical barrier, this study also traces back the dynamics of the Symbiodiniaceae community associated with *P. lobata* over time at the colony scale. More particularly, the Symbiodiniaceae community associated with this *P. lobata* colony is composed of 1 OTU (OTU1) assigned to the *ITS2*-C15-type. This occurrence is largely predominant since the earliest part of the coral core; that was estimated to date back to the 1870’s approximately. Two other Symbiodiniaceae OTUs were found to be (nearly) ubiquitous although at very low proportion even in the contemporaneous living coral (OTU9 and OTU10). These three OTUs are thus likely to be in tight association with the coral host and probably part of the core Symbiodiniaceae community of the *P. lobata* colony. This is in line with previous studies that have documented a similar pattern in the symbiotic community associated with natural colonies of *P. lobata* where the Symbiodiniaceae C15 strain was highly predominant ^43^. No other clear pattern was observed in terms of community changes over time except for the most abundant non-ubiquitous OTUs such as OTU2, OTU4 and OTU5 which are generally absent in the oldest *cora*DNA samples and appear sporadically only from the S06 – S08 samples (i.e. since the 1920’s) and upward along the core. Two hypotheses could explain this temporal fluctuation of these OTUs. First, the abundance of these OTUs in the environment display important temporal fluctuations and the studied *P. lobata* colony adjusted its associated Symbiodiniaceae community according to the most prevalent OTU present in the environment. Alternatively, the abundance of the different Symbiodiniaceae strains in the environment is relatively stable in time and the *P. lobata* colony adjusted its Symbiodiniaceae community according to its interaction’ preference, physiological state and/or to the prevailing environment ^44^. Importantly however, and considering that each OTU is a singular genetic entity, we lack information regarding the functional and physiological consequence of the association between these different *Cladocopium* strains and *P. lobata* to support one of these hypotheses. The use of more resolutive genetic markers could make it possible to better distinguish between different strains from the C15 clade potentially associated with *P. lobata* colonies(ref). Importantly however, and because ancient DNA is more prone to degradation, we could be constrained by the amplicon size. We would thus be more in favor of combining multiple small barcodes rather than using a larger marker.

### Dynamics of Symbiodiniaceae associated with extreme climatic events

No major changes in the Symbiodiniaceae community were observed after the targeted severe ENSO event that occurred in 1997-1998 and which led to a decrease in growth visible on the core, compared to that observed in the *cora*DNA sampled before this event. Two hypotheses might explain this result. First, the targeted ENSO event did not stress the colony enough (e.g. no thermal bleaching occurred) to temporally modify the associated Symbiodiniaceae community. In this regard, and although not comparable to a bleaching event, stability of the Symbiodiniaceae community associated with *P. lobata* during extreme storm events was previously reported on Kiritimati Island in the central equatorial Pacific Ocean ^34^. More generally, the association between *P. lobata* and their symbionts are generally acknowledged to be stable over time even when facing environmental stress ^34,43^. Alternatively, and despite all precautions, we might have missed the succession corresponding to the past coral colony recovering from the ENSO event during the *cora*DNA sampling. Clearly, more coraDNA replicates could have allowed teasing apart these two hypotheses. However, because of the difficulty of sampling only a single succession of coral growth band, and the limited amount of material that can be obtained, replicating coraDNA samples is particularly tedious. One solution could be to use a drill to excavate larger-diameter cores from natural colonies. The most important pattern observed in this study however, concerns the apparent loss of the Symbiodiniaceae community diversity observed post 2010 abnormal warm winter period ^27^. In fact, the E04 *cora*DNA sample harbor a very different symbiotic community compared to the 3 others recent *cora*DNA samples and to the control. Only 3 Symbiodiniaceae OTUs were detected in this *cora*DNA sample while 8 to 10 were found in the other 3 recent samples and in the contemporaneous control sample. Such low diversity found in E04 cannot be explained by a lower sequencing coverage (Supplementary Figure SF1). One hypothesis could be that this pattern reflects a temporary and partial loss of Symbiodiniaceae diversity by the coral colony during this abnormally warm and stable period over the year. This is in line with a pattern of decreased in symbiotic algae diversity observed in less variable environments in several coral species ^45^. Moreover, the remaining 3 OTUs detected in E04 do not correspond to the most abundant ones, which suggest that this loss in *Symbiodinium* OTUs is likely to be non-random. These 3 OTUs are those that are ubiquitous to all *cora*DNA samples. This result hence supports the hypothesis that these 3 OTUs are part of the core Symbiodiniaceae community of the *P. lobata* colony irrespective to their (sometimes low) relative abundance among the colony ^15^.

### Conclusion and perspectives

Together our results highlight the fact that ancient DNA can be extracted from cores excavated from stony corals, the so-called *cora*DNA, dating back to at least one century. Moreover, among the DNA material extracted from the core, we detected DNA from the coral host species, the dinoflagellate symbionts and a free-living environmental dinoflagellate. This result paves the way for studying the temporal evolutionary dynamics of coral holobionts at the colony scale. Combined with some geochemical analyses on the same samples collected along the coral core, this approach could provide new insights on the mechanisms underlying the adaptive responses of corals to several past stress events including temperature changes, pH and/or chemicals such as heavy metals.

## Supplementary material

**Supplementary table S1:** Pairwise genetic distances computed between each pair of the sequences from the OTUs obtained in this study or from previously identified *Symbiodinium* strains available from LaJeunesse *et al*. (2003).

|  | OTU_1 | OTU_2 | OTU_3 | OTU_4 | OTU_5 | OTU_6 | OTU_8 | OTU_9 | OTU_10 |
| --- | --- | --- | --- | --- | --- | --- | --- | --- | --- |
| OTU_1 | 0 | 0,014 | 0 | 0,007 | 0,007 | 0 | 0,044 | 0,022 | 0,022 |
| OTU_2 | 0,014 | 0 | 0,014 | 0,022 | 0,022 | 0,014 | 0,059 | 0,036 | 0,036 |
| OTU_3 | 0 | 0,014 | 0 | 0,007 | 0,007 | 0 | 0,044 | 0,022 | 0,022 |
| OTU_4 | 0,007 | 0,022 | 0,007 | 0 | 0,014 | 0,007 | 0,051 | 0,029 | 0,029 |
| OTU_5 | 0,007 | 0,022 | 0,007 | 0,014 | 0 | 0,007 | 0,051 | 0,029 | 0,029 |
| OTU_6 | 0 | 0,014 | 0 | 0,007 | 0,007 | 0 | 0,044 | 0,022 | 0,022 |
| OTU_8 | 0,044 | 0,059 | 0,044 | 0,051 | 0,051 | 0,044 | 0 | 0,067 | 0,067 |
| OTU_9 | 0,022 | 0,036 | 0,022 | 0,029 | 0,029 | 0,022 | 0,067 | 0 | 0,029 |
| OTU_10 | 0,022 | 0,036 | 0,022 | 0,029 | 0,029 | 0,022 | 0,067 | 0,029 | 0 |
| AY239363.1_Symb_C1b | 0,029 | 0,044 | 0,029 | 0,037 | 0,037 | 0,029 | 0,075 | 0,051 | 0,051 |
| AY239364.1_Symb_C1c | 0,029 | 0,044 | 0,029 | 0,037 | 0,037 | 0,029 | 0,075 | 0,051 | 0,051 |
| AY239365.1_Symb_C3h | 0,022 | 0,036 | 0,022 | 0,029 | 0,029 | 0,022 | 0,067 | 0,044 | 0,044 |
| AY239366.1_Symb_C3j | 0,022 | 0,036 | 0,022 | 0,029 | 0,029 | 0,022 | 0,067 | 0,044 | 0,044 |
| AY239367.1_Symb_C8 | 0,037 | 0,052 | 0,037 | 0,044 | 0,044 | 0,037 | 0,083 | 0,059 | 0,059 |
| AY239368.1_Symb_C8a | 0,044 | 0,059 | 0,044 | 0,052 | 0,052 | 0,044 | 0,091 | 0,067 | 0,067 |
| AY239369.1_Symb_C15 | 0 | 0,014 | 0 | 0,007 | 0,007 | 0 | 0,044 | 0,022 | 0,022 |
| AY239370.1_Symb_C17 | 0,022 | 0,036 | 0,022 | 0,029 | 0,029 | 0,022 | 0,067 | 0,044 | 0,044 |
| AY239372.1_Symb_C21 | 0,022 | 0,036 | 0,022 | 0,029 | 0,029 | 0,022 | 0,067 | 0,044 | 0,044 |
| AY239378.1_Symb_C26 | 0,022 | 0,036 | 0,022 | 0,029 | 0,029 | 0,022 | 0,067 | 0,044 | 0,044 |
| AY258488.1_Symb_C1d | 0,036 | 0,051 | 0,036 | 0,044 | 0,044 | 0,036 | 0,082 | 0,059 | 0,059 |
| AY258496.1_Symb_C31 | 0,029 | 0,044 | 0,029 | 0,037 | 0,037 | 0,029 | 0,075 | 0,051 | 0,051 |
| AY258501.1_Symb_C35a | 0,022 | 0,036 | 0,022 | 0,029 | 0,029 | 0,022 | 0,067 | 0,044 | 0,044 |
| AY765398.1_Symb_C1m | 0,037 | 0,052 | 0,037 | 0,044 | 0,044 | 0,037 | 0,083 | 0,059 | 0,059 |
| AY765400.2_Symb_C33 | 0,044 | 0,059 | 0,044 | 0,052 | 0,052 | 0,044 | 0,091 | 0,067 | 0,067 |
| AY765401.1_Symb_C35 | 0,036 | 0,051 | 0,036 | 0,044 | 0,044 | 0,036 | 0,082 | 0,059 | 0,059 |
| AY765402.1_Symb_C42 | 0,029 | 0,044 | 0,029 | 0,037 | 0,037 | 0,029 | 0,075 | 0,051 | 0,051 |
| AY765413.1_Symb_C78 | 0,036 | 0,051 | 0,036 | 0,044 | 0,044 | 0,036 | 0,082 | 0,059 | 0,059 |
| AY765414.2_Symb_C79 | 0,044 | 0,059 | 0,044 | 0,052 | 0,052 | 0,044 | 0,091 | 0,067 | 0,067 |
| AF333518.1_Symb_C4 | 0,014 | 0,029 | 0,014 | 0,022 | 0,022 | 0,014 | 0,059 | 0,036 | 0,036 |
| AY589730.1_Symb_C1g | 0,029 | 0,044 | 0,029 | 0,037 | 0,037 | 0,029 | 0,075 | 0,051 | 0,051 |
| AY589734.1_Symb_C3f | 0,029 | 0,044 | 0,029 | 0,036 | 0,036 | 0,029 | 0,075 | 0,051 | 0,051 |
| AY589746.1_Symb_C31a | 0,029 | 0,044 | 0,029 | 0,037 | 0,037 | 0,029 | 0,075 | 0,051 | 0,051 |
| AY589752.1_Symb_C47 | 0,029 | 0,044 | 0,029 | 0,037 | 0,037 | 0,029 | 0,075 | 0,051 | 0,051 |
| AY589771.1_Symb_C66 | 0,029 | 0,044 | 0,029 | 0,037 | 0,037 | 0,029 | 0,075 | 0,051 | 0,051 |
| AY258473.1_Symb_C1h | 0,029 | 0,044 | 0,029 | 0,037 | 0,037 | 0,029 | 0,075 | 0,051 | 0,051 |
| AY258475.1_Symb_C8b | 0,037 | 0,052 | 0,037 | 0,044 | 0,044 | 0,037 | 0,083 | 0,059 | 0,059 |
| AY258481.1_Symb_C37 | 0,052 | 0,067 | 0,052 | 0,06 | 0,06 | 0,052 | 0,099 | 0,075 | 0,075 |
| AY258485.1_Symb_C40 | 0,036 | 0,051 | 0,036 | 0,044 | 0,044 | 0,036 | 0,082 | 0,059 | 0,059 |
| AY258487.1_Symb_C42 | 0,036 | 0,051 | 0,036 | 0,044 | 0,044 | 0,036 | 0,082 | 0,059 | 0,059 |
| AF499789.1_Symb_C3 | 0,022 | 0,036 | 0,022 | 0,029 | 0,029 | 0,022 | 0,067 | 0,044 | 0,044 |
| AF499794.1_Symb_C4 | 0,029 | 0,044 | 0,029 | 0,037 | 0,037 | 0,029 | 0,075 | 0,051 | 0,051 |
| AF499797.1_Symb_C7 | 0,022 | 0,036 | 0,022 | 0,029 | 0,029 | 0,022 | 0,067 | 0,044 | 0,044 |
| AF499801.1_Symb_C12 | 0,022 | 0,036 | 0,022 | 0,029 | 0,029 | 0,022 | 0,067 | 0,044 | 0,044 |
| JQ003816.1_Symb_D | 0,827 | 0,82 | 0,827 | 0,856 | 0,827 | 0,827 | 0,856 | 0,82 | 0,861 |

**Supplementary table S2:** Raw data of the sequence abundance of each OTU identified in this study over all *cora*DNA samples.

| OTU_<br>id | Nseq<br>total | S01 | S04 | S04" | S05 | S06 | S08 | S08" | S09 | S10 | S11 | S12 | S13 | S14 | S15 | E01 |
| --- | --- | --- | --- | --- | --- | --- | --- | --- | --- | --- | --- | --- | --- | --- | --- | --- |
| OTU_1 | 748108 | 55576 | 44178 | 12986 | 3924 | 39658 | 62753 | 43406 | 37033 | 3493 | 41942 | 1476 | 67729 | 68005 | 39137 | 67107 |
| OTU_2 | 32296 | 0 | 1 | 0 | 0 | 2912 | 2786 | 340 | 6099 | 0 | 2030 | 0 | 4994 | 3 | 9241 | 1460 |
| OTU_3 | 23547 | 8773 | 0 | 0 | 0 | 0 | 3858 | 172 | 0 | 0 | 694 | 0 | 0 | 0 | 0 | 4078 |
| OTU_4 | 6503 | 0 | 0 | 0 | 0 | 0 | 1574 | 84 | 0 | 0 | 553 | 0 | 3653 | 0 | 0 | 237 |
| OTU_5 | 1193 | 0 | 0 | 0 | 0 | 0 | 0 | 333 | 0 | 386 | 0 | 0 | 0 | 0 | 0 | 225 |
| OTU_6 | 264 | 0 | 0 | 0 | 0 | 0 | 0 | 0 | 0 | 0 | 0 | 0 | 0 | 0 | 0 | 123 |
| OTU_7 | 230 | 0 | 0 | 0 | 0 | 0 | 211 | 19 | 0 | 0 | 0 | 0 | 0 | 0 | 0 | 0 |
| OTU_8 | 141 | 0 | 0 | 0 | 0 | 0 | 113 | 0 | 0 | 0 | 0 | 0 | 0 | 0 | 0 | 0 |
| OTU_9 | 139 | 8 | 11 | 5 | 1 | 6 | 10 | 1 | 9 | 2 | 9 | 0 | 15 | 15 | 6 | 7 |
| OTU_10 | 114 | 10 | 13 | 2 | 3 | 7 | 8 | 8 | 6 | 3 | 8 | 0 | 12 | 8 | 4 | 4 |
| OTU_11 | 92 | 81 | 0 | 0 | 0 | 0 | 4 | 0 | 0 | 0 | 0 | 0 | 0 | 0 | 0 | 0 |
| OTU_12 | 94 | 0 | 0 | 0 | 0 | 0 | 0 | 94 | 0 | 0 | 0 | 0 | 0 | 0 | 0 | 0 |
| OTU_13 | 93 | 0 | 0 | 0 | 0 | 0 | 49 | 44 | 0 | 0 | 0 | 0 | 0 | 0 | 0 | 0 |

SF1: Rarefaction curves obtained for each of the 19 samples.

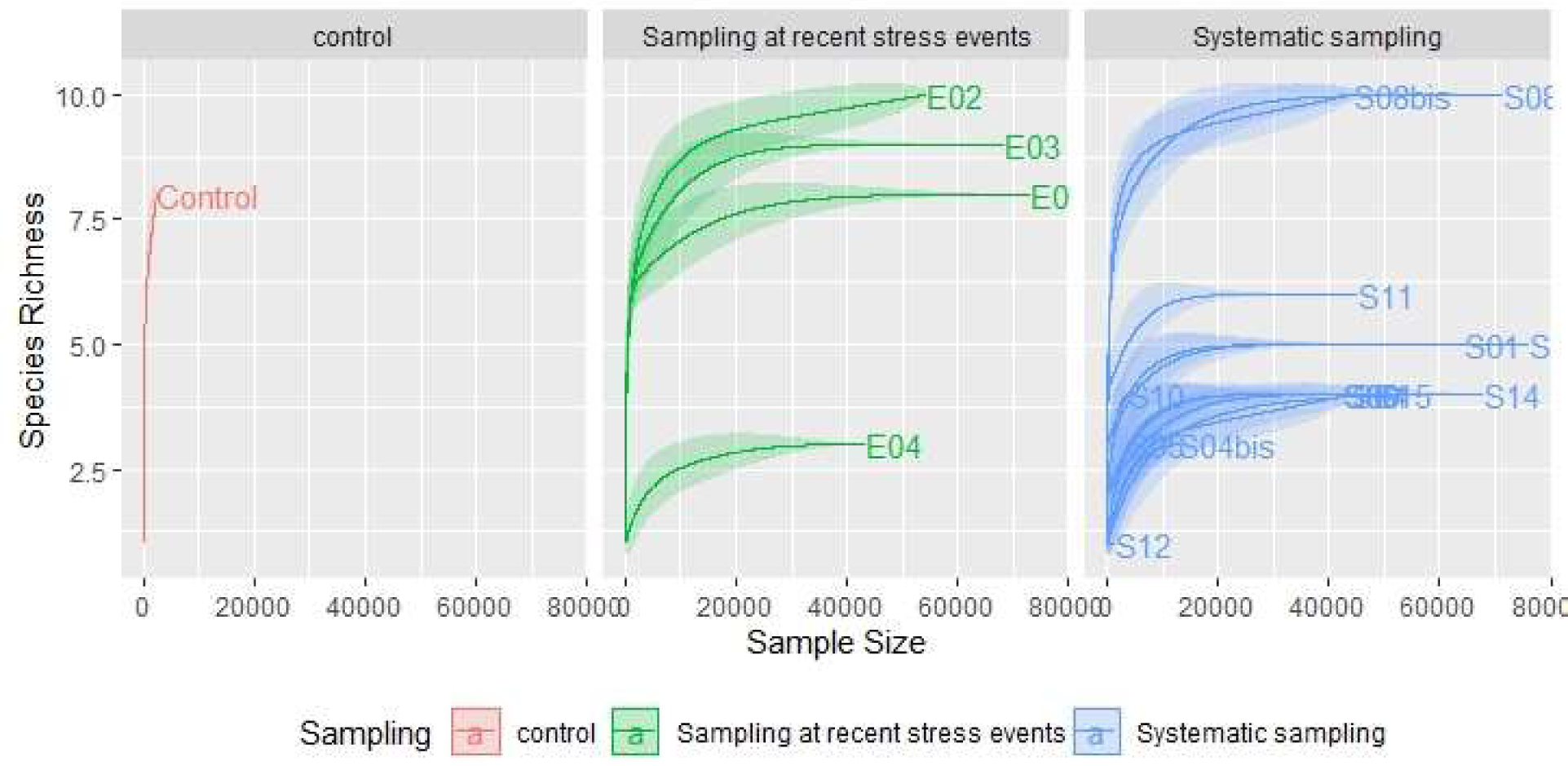

## Declarations

### Availability of data and materials

Raw sequences from the metabarcoding approach are available on GenBank under the SRA project id. PRJNA1086717.

### Authors’ contributions

O.R and J.V.D equally contributed to the conception and design of the work. The coral core was sampled and sent to the IHPE laboratory by D.D and J.B with the help of G.I for administrative aspects. D.D. and T.G. dated the coral core successions. T.G, M.S, O.R, M.B.Z and J.V.D acquired the ancient DNA extracts with the help of C.T at the degraded DNA platform (Montpellier). O.R, J.F, M.S and J.V.D processed the DNA extracts. O.R., T.G and E.T. analyzed the metabarcoding datasets. O.R., T.G. and J.V.D interpreted the results. O.R have drafted the manuscript and J.V.D, D.D and E.T. substantively revised it. All authors approved the submitted version.

## Acknowledgments

O.R and J.V.D would like to thank Diane Merceron for her kind help in getting the coral core travelling from New Caledonia to Perpignan. We thank the personal (divers, boat pilot, etc,..) of the IRD centre of New Caledonia who allowed for coral core extraction and transportation. We thank the Bio-Environment platform (University of Perpignan Via Domitia) for the sequencing process, Dr. Cristian Chaparro and the IHPE bioinformatics service and the GenoToul bioinformatics platform, Toulouse Occitanie (Bioinfo Genotoul, doi: 10.15454/1.5572369328961167E12) for providing computing resources (Galaxy instance; http://sigenae-workbench.toulouse.inra.fr). Finally, we would like to warmly thanks all staff from the imagery service at the public Hospital of Perpignan for their kind welcome and help during CT scan including Stephane Belfio, Marine Maurel and Cécilia Colomer. This study is set within the framework of the “Laboratoire d’Excellence (LabEx)” TULIP (ANR-10-LABX-41) and with the technical support of the LabEx CeMEB (ANR-10-LABX-04-01).

## References

1. Díaz, S., Settele, J., Brondízio, E., Ngo, H.T., Guèze, M., Agard, J., Arneth, A., Balvanera, P., Brauman, K., Watson, R.T., et al. (2019). Résumé à l’intention des décideurs du rapport sur l’évaluation mondiale de la biodiversité et des services écosystémiques de la Plateforme intergouvernementale scientifique et politique sur la biodiversité et les services écosystémiques. 53.

2. DeCarlo, T.M., Harrison, H.B., Gajdzik, L., Alaguarda, D., Rodolfo-Metalpa, R., D’Olivo, J., Liu, G., Patalwala, D., and McCulloch, M.T. (2019). Acclimatization of massive reef-building corals to consecutive heatwaves. Proc. R. Soc. B 286, 20190235. 10.1098/rspb.2019.0235.

3. Maggioni, F., Pujo-Pay, M., Aucan, J., Cerrano, C., Calcinai, B., Payri, C., Benzoni, F., Letourneur, Y., and Rodolfo-Metalpa, R. (2021). The Bouraké semi-enclosed lagoon (New Caledonia) – a natural laboratory to study the lifelong adaptation of a coral reef ecosystem to extreme environmental conditions. Biogeosciences 18, 5117–5140. 10.5194/bg-18-5117-2021.

4. Palumbi, S.R., Barshis, D.J., Traylor-Knowles, N., and Bay, R.A. (2014). Mechanisms of reef coral resistance to future climate change. Science 344, 895–898. 10.1126/science.1251336.

5. Clerissi, C., Brunet, S., Vidal-Dupiol, J., Adjeroud, M., Lepage, P., Guillou, L., Escoubas, J.-M., and Toulza, E. (2018). Protists Within Corals: The Hidden Diversity. Front. Microbiol. 9, 2043. 10.3389/fmicb.2018.02043.

6. Webster, N.S., and Reusch, T.B.H. (2017). Microbial contributions to the persistence of coral reefs. ISME J 11, 2167–2174. 10.1038/ismej.2017.66.

7. Dixon, G.B., Davies, S.W., Aglyamova, G.V., Meyer, E., Bay, L.K., and Matz, M.V. (2015). Genomic determinants of coral heat tolerance across latitudes. Science 348, 1460– 1462. 10.1126/science.1261224.

8. Van Oppen, M.J.H., Souter, P., Howells, E.J., Heyward, A., and Berkelmans, R. (2011). Novel Genetic Diversity Through Somatic Mutations: Fuel for Adaptation of Reef Corals? Diversity 3, 405–423. 10.3390/d3030405.

9. Rey, O., Danchin, E., Mirouze, M., Loot, C., and Blanchet, S. (2016). Adaptation to Global Change: A Transposable Element–Epigenetics Perspective. Trends in Ecology & Evolution 31, 514–526. 10.1016/j.tree.2016.03.013.

10. Rinkevich, B. (2019). Coral chimerism as an evolutionary rescue mechanism to mitigate global climate change impacts. Glob Change Biol 25, 1198–1206. 10.1111/gcb.14576.

11. Vidal-Dupiol, J., Harscouet, E., Shefy, D., Toulza, E., Rey, O., Allienne, J.-F., Mitta, G., and Rinkevich, B. (2022). Frontloading of stress response genes enhances robustness to environmental change in chimeric corals. BMC Biol 20, 167. 10.1186/s12915-022-01371-7.

12. Liew, Y.J., Howells, E.J., Wang, X., Michell, C.T., Burt, J.A., Idaghdour, Y., and Aranda, M. (2020). Intergenerational epigenetic inheritance in reef-building corals. Nat. Clim. Chang. 10, 254–259. 10.1038/s41558-019-0687-2.

13. Torda, G., Donelson, J.M., Aranda, M., Barshis, D.J., Bay, L., Berumen, M.L., Bourne, D.G., Cantin, N., Foret, S., Matz, M., et al. (2017). Rapid adaptive responses to climate change in corals. Nature Climate Change 7, 627–636. 10.1038/nclimate3374.

14. Berkelmans, R., and van Oppen, M.J.H. (2006). The role of zooxanthellae in the thermal tolerance of corals: a ‘nugget of hope’ for coral reefs in an era of climate change. Proc. R. Soc. B 273, 2305–2312. 10.1098/rspb.2006.3567.

15. Brener-Raffalli, K., Clerissi, C., Vidal-Dupiol, J., Adjeroud, M., Bonhomme, F., Pratlong, M., Aurelle, D., Mitta, G., and Toulza, E. (2018). Thermal regime and host clade, rather than geography, drive Symbiodinium and bacterial assemblages in the scleractinian coral Pocillopora damicornis sensu lato. Microbiome 6, 39. 10.1186/s40168-018-0423-6.

16. LaJeunesse, T.C., Parkinson, J.E., Gabrielson, P.W., Jeong, H.J., Reimer, J.D., Voolstra, C.R., and Santos, S.R. (2018). Systematic Revision of Symbiodiniaceae Highlights the Antiquity and Diversity of Coral Endosymbionts. Current Biology 28, 2570–2580.e6. 10.1016/j.cub.2018.07.008.

17. Swain, T.D., Chandler, J., Backman, V., and Marcelino, L. (2017). Consensus thermotolerance ranking for 110 Symbiodinium phylotypes: an exemplar utilization of a novel iterative partial-rank aggregation tool with broad application potential. Funct Ecol 31, 172–183. 10.1111/1365-2435.12694.

18. Buerger, P., Alvarez-Roa, C., Coppin, C.W., Pearce, S.L., Chakravarti, L.J., Oakeshott, J.G., Edwards, O.R., and van Oppen, M.J.H. (2020). Heat-evolved microalgal symbionts increase coral bleaching tolerance. Sci. Adv. 6, eaba2498. 10.1126/sciadv.aba2498.

19. Cunning, R., Silverstein, R.N., and Baker, A.C. (2015). Investigating the causes and consequences of symbiont shuffling in a multi-partner reef coral symbiosis under environmental change. Proc. R. Soc. B. 282, 20141725. 10.1098/rspb.2014.1725.

20. Matthews, J.L., Oakley, C.A., Lutz, A., Hillyer, K.E., Roessner, U., Grossman, A.R., Weis, V.M., and Davy, S.K. (2018). Partner switching and metabolic flux in a model cnidarian–dinoflagellate symbiosis. Proc. R. Soc. B. 285, 20182336. 10.1098/rspb.2018.2336.

21. Voolstra, C.R., and Ziegler, M. (2020). Adapting with Microbial Help: Microbiome Flexibility Facilitates Rapid Responses to Environmental Change. BioEssays 42, 2000004. 10.1002/bies.202000004.

22. Hofreiter, M., Paijmans, J.L.A., Goodchild, H., Speller, C.F., Barlow, A., Fortes, G.G., Thomas, J.A., Ludwig, A., and Collins, M.J. (2015). The future of ancient DNA: Technical advances and conceptual shifts: Prospects & Overviews. BioEssays 37, 284–293. 10.1002/bies.201400160.

23. Baker, D.M., Weigt, L., Fogel, M., and Knowlton, N. (2013). Ancient DNA from Coral-Hosted Symbiodinium Reveal a Static Mutualism over the Last 172 Years. PLoS ONE 8, e55057. 10.1371/journal.pone.0055057.

24. DiBattista, J.D., Reimer, J.D., Stat, M., Masucci, G.D., Biondi, P., De Brauwer, M., and Bunce, M. (2019). Digging for DNA at depth: rapid universal metabarcoding surveys (RUMS) as a tool to detect coral reef biodiversity across a depth gradient. PeerJ 7, e6379. 10.7717/peerj.6379.

25. Gomez Cabrera, M., Young, J.M., Roff, G., Staples, T., Ortiz, J.C., Pandolfi, J.M., and Cooper, A. (2019). Broadening the taxonomic scope of coral reef palaeoecological studies using ancient DNA. Mol Ecol 28, 2636–2652. 10.1111/mec.15038.

26. Smith, L.W., Barshis, D., and Birkeland, C. (2007). Phenotypic plasticity for skeletal growth, density and calcification of Porites lobata in response to habitat type. Coral Reefs 26, 559–567. 10.1007/s00338-007-0216-z.

27. Wu, H.C., Dissard, D., Douville, E., Blamart, D., Bordier, L., Tribollet, A., Le Cornec, F., Pons-Branchu, E., Dapoigny, A., and Lazareth, C.E. (2018). Surface ocean pH variations since 1689 CE and recent ocean acidification in the tropical South Pacific. Nat Commun 9, 2543. 10.1038/s41467-018-04922-1.

28. Quigley, K.M., Davies, S.W., Kenkel, C.D., Willis, B.L., Matz, M.V., and Bay, L.K. (2014). Deep-Sequencing Method for Quantifying Background Abundances of Symbiodinium Types: Exploring the Rare Symbiodinium Biosphere in Reef-Building Corals. PLoS ONE 9, e94297. 10.1371/journal.pone.0094297.

29. Lajeunesse, T., and Trench, R. (2000). Biogeography of two species of Symbiodinium (Freudenthal) inhabiting the intertidal sea anemone *Anthopleura elegantissima* (Brandt). The Biological Bulletin 199, 126–134. 10.2307/1542872.

30. Afgan, E., Baker, D., Batut, B., Van Den Beek, M., Bouvier, D., Čech, M., Chilton, J., Clements, D., Coraor, N., and Grüning, B.A. (2018). The Galaxy platform for accessible, reproducible and collaborative biomedical analyses: 2018 update. Nucleic acids research 46, W537–W544.

31. Escudié, F., Auer, L., Bernard, M., Mariadassou, M., Cauquil, L., Vidal, K., Maman, S., Hernandez-Raquet, G., Combes, S., and Pascal, G. (2018). FROGS: Find, Rapidly, OTUs with Galaxy Solution. Bioinformatics 34, 1287–1294. 10.1093/bioinformatics/btx791.

32. LaJeunesse, T.C., Loh, W.K.W., van Woesik, R., Hoegh-Guldberg, O., Schmidt, G.W., and Fitt, W.K. (2003). Low symbiont diversity in southern Great Barrier Reef corals, relative to those of the Caribbean. Limnol. Oceanogr. 48, 2046–2054. 10.4319/lo.2003.48.5.2046.

33. Paradis, E., and Schliep, K. (2019). ape 5.0: an environment for modern phylogenetics and evolutionary analyses in R. Bioinformatics 35, 526–528.

34. Claar, D.C., Tietjen, K.L., Cox, K.D., Gates, R.D., and Baum, J.K. (2020). Chronic disturbance modulates symbiont (Symbiodiniaceae) beta diversity on a coral reef. Sci Rep 10, 4492. 10.1038/s41598-020-60929-z.

35. Thornhill, D.J., Lajeunesse, T.C., and Santos, S.R. (2007). Measuring rDNA diversity in eukaryotic microbial systems: how intragenomic variation, pseudogenes, and PCR artifacts confound biodiversity estimates. Molecular Ecology 16, 5326–5340. 10.1111/j.1365-294X.2007.03576.x.

36. Dabney, J., Meyer, M., and Pääbo, S. (2013). Ancient DNA damage. Cold Spring Harbor perspectives in biology 5, a012567.

37. Salas, R., Tillmann, U., John, U., Kilcoyne, J., Burson, A., Cantwell, C., Hess, P., Jauffrais, T., and Silke, J. (2011). The role of *Azadinium spinosum* (Dinophyceae) in the production of azaspiracid shellfish poisoning in mussels. Harmful Algae 10, 774–783.

38. Akselman, R., and Negri, R.M. (2012). Blooms of *Azadinium cf. spinosum* Elbrächter et Tillmann (Dinophyceae) in northern shelf waters of Argentina, Southwestern Atlantic. Harmful Algae 19, 30–38. 10.1016/j.hal.2012.05.004.

39. Hernández-Becerril, D.U., Barón-Campis, S.A., and Escobar-Morales, S. (2012). A new record of *Azadinium spinosum* (Dinoflagellata) from the tropical Mexican Pacific. Revista de Biología Marina y Oceanografía 47, 553–557.

40. Fu, Z., Piumsomboon, A., Punnarak, P., Uttayarnmanee, P., Leaw, C.P., Lim, P.T., Wang, A., and Gu, H. (2021). Diversity and distribution of harmful microalgae in the Gulf of Thailand assessed by DNA metabarcoding. Harmful Algae 106, 102063.

41. Tillmann, U., Elbrächter, M., Krock, B., John, U., and Cembella, A. (2009). *Azadinium spinosum* gen. et sp. nov.(Dinophyceae) identified as a primary producer of azaspiracid toxins. European Journal of Phycology 44, 63–79.

42. Sibat, M., Mai, T., Tanniou, S., Biegala, I., Hess, P., and Jauffrais, T. (2023). Seasonal Single-Site Sampling Reveals Large Diversity of Marine Algal Toxins in Coastal Waters and Shellfish of New Caledonia (Southwestern Pacific). Toxins 15, 642. 10.3390/toxins15110642.

43. Stat, M., Pochon, X., Franklin, E.C., Bruno, J.F., Casey, K.S., Selig, E.R., and Gates, R.D. (2013). The distribution of the thermally tolerant symbiont lineage (*Symbiodinium* clade D) in corals from Hawaii: correlations with host and the history of ocean thermal stress. Ecol Evol 3, 1317–1329. 10.1002/ece3.556.

44. Baker, A.C. (2003). Flexibility and Specificity in Coral-Algal Symbiosis: Diversity, Ecology, and Biogeography of *Symbiodinium*. Annu. Rev. Ecol. Evol. Syst. 34, 661–689. 10.1146/annurev.ecolsys.34.011802.132417.

45. Thornhill, D.J., LaJeunesse, T.C., Kemp, D.W., Fitt, W.K., and Schmidt, G.W. (2006). Multi-year, seasonal genotypic surveys of coral-algal symbioses reveal prevalent stability or post-bleaching reversion. Marine Biology 148, 711–722. 10.1007/s00227-005-0114-2.

